# Disentangling the fitness cost of gene expression

**DOI:** 10.1101/2025.06.02.656972

**Authors:** Yichen Yan, Jie Lin

**Affiliations:** Peking-Tsinghua Center for Life Sciences, Peking University, Beijing, China; Center for Quantitative Biology, Peking University, Beijing, China; School of Physics, Peking University, Beijing, China

## Abstract

Gene expression is essential for biological functions but also incurs a fitness cost. Although the fitness cost can be experimentally measured as the relative reduction in growth rate, it remains unclear how the cost quantitatively depends on different limiting factors. In this work, we establish a resource competition model and disentangle the fitness cost into components arising from limiting resources, including ribosomes, RNA polymerases, and transcription factors. Comparing our model predictions with experimental data for *Saccharomyces cerevisiae*, we demonstrate that ribosome competition dominates the translation cost, and that transcription factor competition dominates the transcription cost. Our model reveals that the fitness costs originate from the processes of transcription and translation, rather than from the products. The model also systematically connects the fitness cost to genetic and environmental properties, making quantitative predictions consistent with various experimental observations. Our work establishes a systematic framework for gene expression cost, guiding synthetic biology to optimize genetic design.

## INTRODUCTION

Gene expression is one of the most fundamental cellular processes, where genes are transcribed into mRNAs, and mRNAs are translated into proteins critical for a cell’s viability. However, gene expression also imposes a fitness cost: expression of unnecessary proteins burdens the cell, slowing its growth rate [1–7]. The fitness cost is typically quantified by the relative reduction in growth rate [8]. Despite its fundamental importance in the evolution of gene expression, the sources of the fitness cost are not well understood. One potential source is the energy or metabolic cost of gene expression, which arises from the consumption of metabolites, e.g., ATP, nucleotides, and amino acids [5, 9, 10]. Nevertheless, the energy cost fails to explain the fitness cost, as producing mRNA imposes a fitness cost comparable to that of producing proteins [8], even though mRNA synthesis consumes far less ATP than protein synthesis [5, 9]. In addition, because the relationship between numerous biochemical processes and cell growth is complex, predicting fitness cost based on metabolic consumption is challenging.

Prior theories largely attribute the fitness cost to dilution effects, where overexpressed proteins dilute the concentrations of key biomolecules, such as ribosomes and RNA polymerases (RNAPs), leading to a lower growth rate [4, 11]. However, the dilution theory cannot relate the cost to a specific process during gene expression, or to particular properties of the studied gene, which are crucial for synthetic biology. Moreover, the dilution theory disagrees with the experimental observations showing that the gene expression cost is generated by the processes of transcription and translation, rather than by the products; particularly, the fitness cost is independent of the degradation rate of the overexpressed protein [8, 12]. Furthermore, it remains unclear how the fitness cost can be decomposed into a sum of transcription and translation costs [8, 13].

In this work, we calculate the fitness cost based on a resource competition scenario. In the simplest scenario, we assume that genes compete for RNA polymerases for transcription and mRNAs compete for ribosomes for translation. We successfully disentangle the fitness cost into two components generated from the competition for RNAPs and ribosomes. The ribosome cost turns out to be the relative fraction of ribosomes used by the exogenous mRNA, and the RNAP cost is the relative fraction of RNAPs used by the exogenous gene multiplied by the fraction of free ribosomes. Remarkably, our predicted cost due to ribosome competition agrees quantitatively with experimental data for *S. cerevisiae* [8]. Surprisingly, the measured cost generated by transcription is much higher than the RNAP cost.

We next extend our model to include transcription factors (TFs) and show that the cost generated by competition for TFs dominates the transcription cost, and the predicted TF cost aligns with experiments. Our model quantitatively explains why the fitness cost of mRNA production can be comparable to that of protein production despite significantly lower energy expenditure. We also explicitly account for dilution effects and reveal that their net contribution is negligible; therefore, gene expression costs primarily stem from the transcription and translation processes, which offers an alternative to the prevailing assumption that fitness costs arise from the dilution effects of protein products. Our model also quantitatively connects the fitness cost to genetic and environmental properties, showing robust predictive power across diverse biological contexts. From an application perspective, our work establishes a systematic framework for the fitness cost of gene expression, providing design guidelines for synthetic biology and facilitating genetic circuit optimization.

## RESULTS

### Resource competition model of gene expression

We introduce a gene expression model at the genomewide level, in which genes compete for RNA polymerase (RNAP) to transcribe mRNAs, and mRNAs compete for ribosomes to translate proteins (Figure 1) [14, 15]. In eukaryotes, “RNAP” in this work refers to RNA polymerase II, which transcribes mRNAs. We consider the fitness cost of an exogenous gene, denoted by “ex”. To simplify without affecting the main results, we treat all the endogenous genes as an ensemble of “average” genes so that they have the same binding affinity to RNAP and their mRNAs have the same binding affinity to the ribosome. The exogenous gene can have different binding affinities from the “average” ones. One should note that our results are equally valid for an overexpressed endogenous gene, and one only needs to treat the additional copy of the endogenous gene as the “exogenous” gene.

**FIG. 1.**
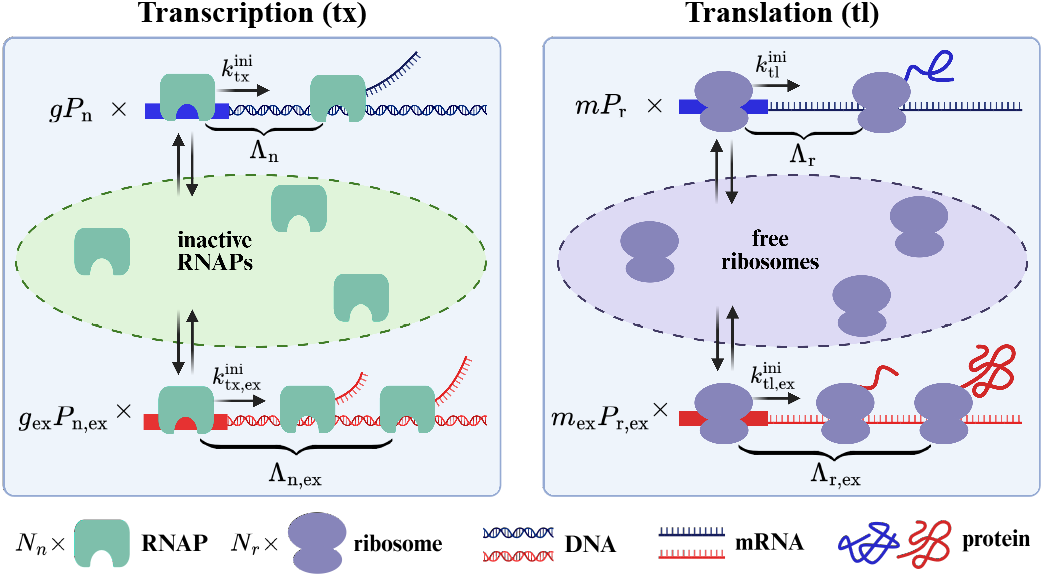
Schematic of the resource competition model. The exogenous gene and the endogenous genes compete for the limiting RNAP and ribosomes. RNAP can be actively transcribing endogenous genes, actively transcribing the exogenous gene, or inactive. Their numbers sum to a fixed total RNAP copy number. Endogenous genes (blue) and the exogenous gene compete for RNAP, where *g* and *g*_ex_ are their copy numbers, and *P*_n_ and *P*_n,ex_ are their probabilities for promoters to be bound by an RNAP, respectively. Similarly, endogenous mRNAs (blue) and the exogenous mRNAs (red) compete for ribosomes, where *m* and *m*_ex_ are their copy numbers, and *P*_r_ and *P*_r,ex_ are probabilities for their RBSs to be bound by a ribosome, respectively.

At the transcription level, RNAP can be in three states (Figure 1): actively transcribing endogenous genes, actively transcribing the exogenous gene, and inactive (i.e., free or bound to non-coding regions). Without the exogenous gene, the conservation of the total RNAP copy number can be written as

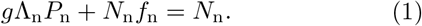

Here, *g* is the genome size, i.e., the total copy number of endogenous genes; Λ_n_ is the gene capacity of RNAP, i.e., the maximum number of RNAP on a single gene copy, including initiating and elongating RNAP; *P*_n_ is the probability of an endogenous promoter bound by an RNAP, which we model as a Michaelis-Menten (MM) function of the concentration of free RNAP [16–18] (Methods A); *N*_n_ is the total number of RNAP; and *f*_n_ is the fraction of inactive RNAP molecules which can be either free or bound to non-coding regions.

The introduction of the exogenous gene leads to the changes in *P*_n_ and *f*_n_: Δ*P*_n_ and Δ*f*_n_, which can be found by

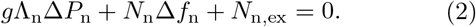

Here, *N*_n,ex_ is the number of RNAP working on the exogenous gene. In this work, we mainly consider a simplified scenario in which the resource pool is fixed (e.g., Δ*N*_n_ = 0), and we later show that this is a good approximation because the effects of changes in the resource copy number and the cell volume are largely canceled out. Eq. (2) states that the resource competition from the exogenous gene leads to a reduction in both the inactive RNAP fraction and the probability of endogenous promoters bound by an RNAP.

The equilibrium of production and degradation sets the total endogenous mRNA copy number:

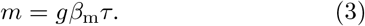

Here, 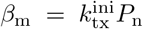 is the mRNA production rate per gene, where 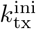is the transcription initiation rate for an RNAP-bound promoter (number of initiations per unit time), and *τ* is the mRNA lifetime.

Without the exogenous gene, a similar conservation equation applies to ribosomes:

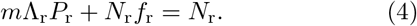

Here, *m* is the copy number of endogenous mRNA; Λ_r_ is the mRNA capacity of ribosomes (i.e., the maximum number of ribosomes on a single mRNA copy); *P*_r_ is the probability for the ribosome-binding-site (RBS) of endogenous mRNA to be bound by a ribosome, which we model as an MM function of the free ribosome concentration [15, 19, 20] (Methods A); *N*_r_ is the total number of ribosomes; *f*_r_ is the fraction of free ribosomes that are not actively involved in translation.

The introduction of the exogenous gene leads to changes in *m, P*_r_, and *f*_r_: Δ*m*, Δ*P*_r_, and Δ*f*_r_, which can be found by

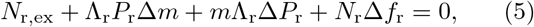

where *N*_r,ex_ is the number of ribosomes working on the exogenous mRNA produced by the exogenous gene. We note that the change in the endogenous mRNA copy number originates from a lower mRNA production rate due to the reduced *P*_n_ after introducing the exogenous gene (Eqs. (2-3)).

The time derivative of endogenous protein mass *M* is set by production and degradation:

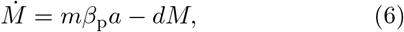

where *a* is the mass of the “average” endogenous protein, and *d* is the average protein degradation rate. Here, 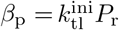 is the protein production rate per mRNA, where 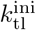 is the translation initiation rate for a ribosome-bound RBS. Because the growth rates of all extensive variables are the same in the exponential steady state, we calculate the growth rate using the endogenous protein mass as

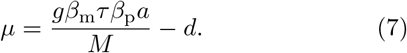

### Decomposing the fitness cost of gene expression

We introduce the fitness cost defined as the relative reduction in the growth rate due to the expression of the exogenous gene, assuming the exogenous gene generates no benefit:

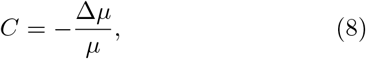

Intriguingly, the resource competition model predicts that the fitness cost of gene expression can be explicitly disentangled into a cost due to RNAP competition (RNAP cost) and a cost due to ribosome competition (ribosome cost) (Figure 2; derivation details in Methods A):

**FIG. 2.**
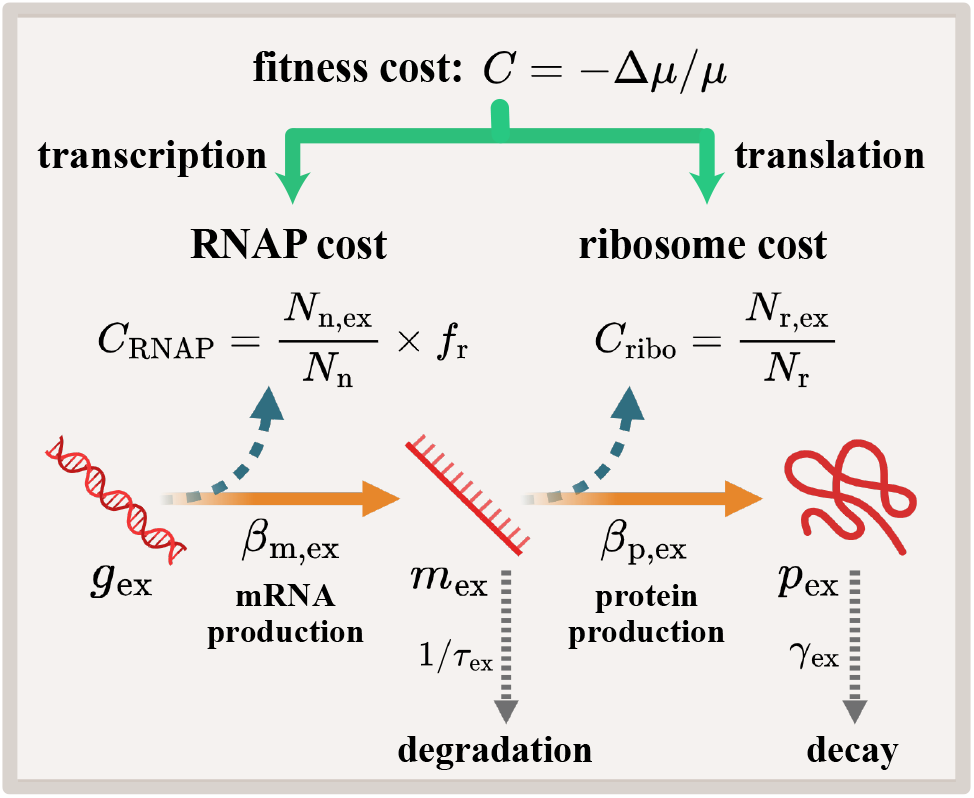
The RNAP and ribosome costs of gene expression. The fitness cost can be disentangled into a ribosome cost and an RNAP cost. The ribosome cost equals the fraction of ribosomes used by the exogenous mRNA (*N*_r,ex_*/N*_r_). The RNAP cost equals the fraction of RNAPs used by the exogenous gene (*N*_n,ex_*/N*_n_) multiplied by the free ribosome fraction. *β*_m,ex_, *β*_p,ex_, 1*/τ*_ex_, *γ*_ex_ are the rates of mRNA production, protein production, mRNA degradation, and protein decay for the corresponding molecule of the exogenous gene, respectively.

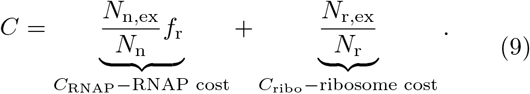

Here, *N*_n,ex_ and *N*_r,ex_ are the copy numbers of RNAP and ribosomes working on the exogenous gene and mRNA, respectively. The above equation shows that the cost of expressing the exogenous gene is determined by the number of RNAPs and ribosomes it consumes. Interestingly, the RNAP cost is damped by a downstream factor *f*_r_, the fraction of free ribosomes in the total ribosome pool: the upstream RNAP cost only becomes apparent when ribosome availability downstream is not limiting.

We can also express the fitness cost as a function of the produced mRNA and protein copy number of the exogenous gene (Methods A):

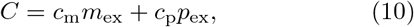

where

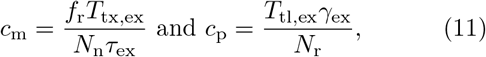

are the RNAP cost per mRNA and the ribosome cost per protein, respectively. Here, *T*_tx,ex_ is the total duration for an RNAP to transcribe the exogenous gene, including the duration in the initiation state. *T*_tl,ex_ is the total duration for a ribosome to translate the corresponding mRNA, including the duration in the initiation state. *τ*_ex_ is the lifetime of the exogenous mRNA, and *γ*_ex_ = *µ* +*d*_ex_ is the decay rate of the exogenous protein, which is the sum of the growth rate and the degradation rate *d*_ex_. Eq. (10) suggests that cells can lower the gene expression cost by using fewer mRNA copies to produce a given amount of proteins, in agreement with experiments [21, 22].

### Limiting transcription factors dominate transcription cost in *S. cerevisiae*

Having established that the fitness cost can be decomposed into a linear combination of RNAP cost and ribosome cost, we analyze experimental data and compare them with our model. Kafri et al. integrated exogenous mCherry genes into *S. cerevisiae* and measured the relative reduction in growth rate due to exogenous gene expression [8]. To compare with their experiments, we estimate the cost coefficients using the resource competition model with the relevant parameters of *S. cerevisiae*.

Interestingly, the predicted ribosome cost is very close to the measured translation cost (Table I), meaning that ribosome competition is the dominant source of translation cost, consistent with previous findings [19, 42, 43].

**TABLE I.** Comparisons between the costs estimated from the experimental data and the resource competition model for an mCherry gene in *S. cerevisiae*. Here, we analyze the data of TDH3 promoter in the standard culture from Ref. [8]. In this table, all variables correspond to one exogenous gene copy (*g*_ex_ = 1) for simplicity. Numerical values are rounded to one significant figure, considering the uncertainties in experiments and estimations. Details of the experimental values are in Methods B. Details of the theoretical estimations are as follows: (1) To estimate 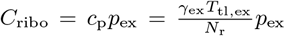 (Eqs. [9-11]), we take the following parameters: the protein decay rate *γ*_ex_ *≈ µ* = 0.4/h [8], the ribosome copy number *N*_r_ = 3 ×10^5^ [23, 24], and the mCherry protein copy number for each gene copy *p*_ex_ = 1.5 ×10^6^ (Methods B). The protein decay rate is dominated by growth rate (*γ*_ex_ ≈*µ*) because degradation is much slower compared to cell growth [25–27]. The duration for a ribosome to translate an mCherry mRNA is estimated as 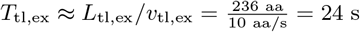, where *L*_tl,ex_ is the length of the exogenous mCherry protein in amino acids [28], and *v*_tl,ex_ is the translation elongation speed of the ribosome [29, 30]. (2) To estimate 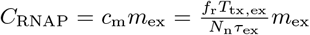 (Eqs. [9-11]), we take the following parameters: the fraction of free ribosomes *f*_r_ = 0.3 [31], the copy number of RNAP *N*_n_ = 3 × 10^4^ [32], the mRNA lifetime *τ*_ex_ = 15 min [33–35], and the mCherry mRNA copy number for each gene copy *m*_ex_ = 6 × 10^2^ (Methods B). The duration for an RNAP to transcribe an mCherry gene is estimated as 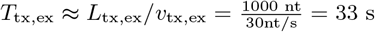, where *L*_tx,ex_ is the length of the exogenous gene transcribed in nucleotides [8], and *v*_tx,ex_ is the transcription elongation speed of RNAP [36–38]. In both (1) and (2), we neglect the duration of initiation, which is typically much shorter than the duration of elongation. (3) An mCherry gene driven by the TDH3 promoter accounts for 3.4% of the total protein number, 200 times more than the contribution of an average gene, which we estimate as 1*/g* with *g* = 6000 [39, 40]. Assuming the primary difference comes from the promoter strength, we take *β*_m,ex_*/β*_m_ = 200. Since the actual fraction of free transcription factors is unknown, we take *f*_t_ = 0.5, which does not affect the order of magnitude. We use Eq. (13) to estimate *c*_g_, which is equal to *C*_TF_ given *g*_ex_ = 1. Other parameters are taken as *f*_n_ = 0.9 [41] and *f*_r_ = 0.3 [31]. We remark that the inactive RNAP fraction *f*_n_ includes both free RNAPs and RNAPs that are associated with non-coding regions. (4) The translation cost *C*_tl_ (theory) is equal to the ribosome cost *C*_ribo_. (5) The transcription cost *C*_tx_ (theory) is the sum of the RNAP cost and the TF cost: *C*_tx_ = *C*_RNAP_ + *C*_TF_.

| Variable (per gene copy) | $C_{\text{ribo}}$ | $C_{\text{RNAP}}$ | $C_{\text{TF}}$ | $C_{\text{tl}}$ | $C_{\text{tx}}$ |
| --- | --- | --- | --- | --- | --- |
| Theory | $1 \times 10^{-2}$ | $2 \times 10^{-4}$ | $5 \times 10^{-3}$ | $1 \times 10^{-2}$ | $5 \times 10^{-3}$ |
| Experiment | — | — | — | $9 \times 10^{-3}$ | $6 \times 10^{-3}$ |

The ratio of the RNAP cost to the ribosome cost is about 1/50 (Table I), comparable to the transcription-translation cost ratio estimated by the ATP consumption rate [5, 9], suggesting that the cellular resources for transcription and translation are coordinated with their energy expenditures. However, the experimentally measured transcription cost is on the same order of magnitude as the translation cost, much higher than the RNAP cost (Table I), which cannot be accounted for by either ATP consumption or RNAP competition. These results suggest that other limiting factors must exist at the transcription level besides RNAP.

Experiments have shown that protein-burdened cells resemble mutants lacking transcription coactivators, including Mediator, SAGA, and SWI-SNF, both transcriptionally and phenotypically [44]. These results suggest that transcription-initiation-associated proteins may also be limiting factors for transcription. Therefore, we extend our model to include the resource competition for transcription-initiation-associated proteins, including general transcription factors, mediators, and chromatin remodeling proteins [45–47]. For simplicity, we combine all these proteins into a coarse-grained type of TF, and the transcription initiation requires the binding of TFs near the promoter (Figure 3a). Therefore, the mRNA production rate also depends on the TF concentration: 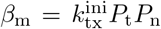 where *P*_t_ is the TF binding probability, which we model as a MM function of the free TF concentration (Methods C). Notably, the fitness cost can be explicitly decomposed as (Methods C)

**FIG. 3.**
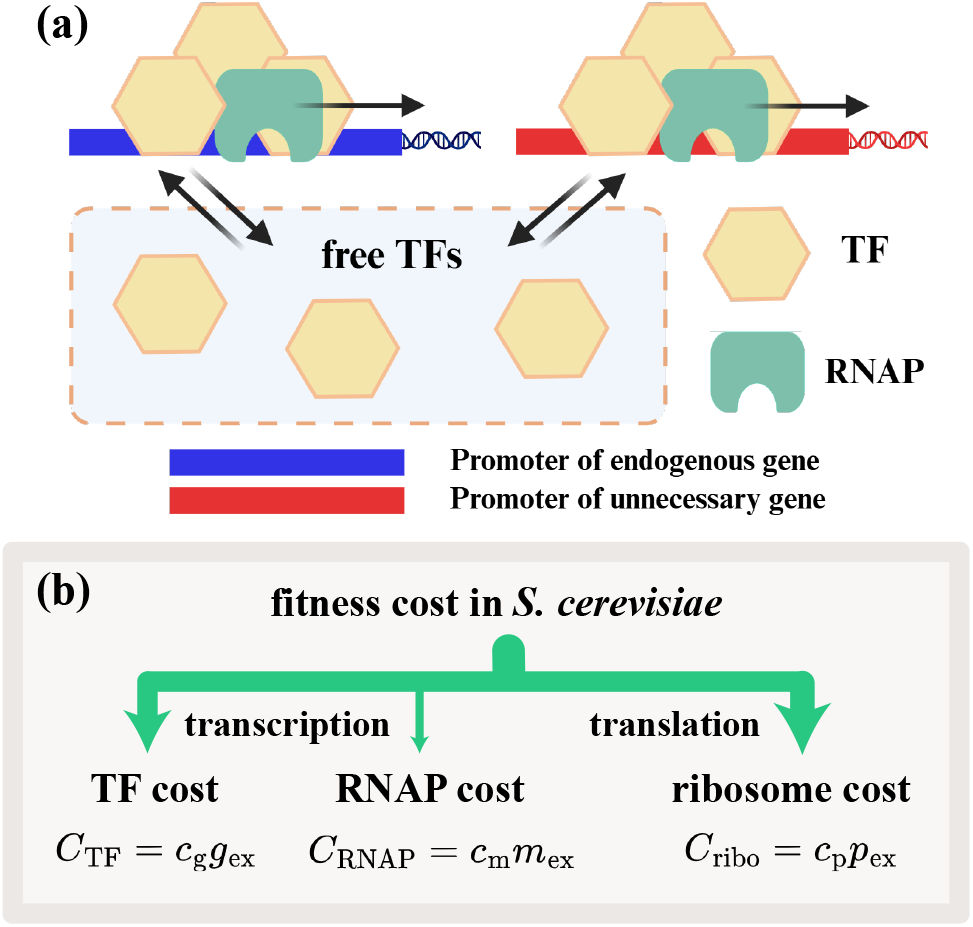
The extended model incorporating the TF cost. (a) Competition for the limiting TFs generates an additional component of transcription cost. (b) The fitness cost in *S. cerevisiae* can be decomposed into a TF cost, an RNAP cost, and a ribosome cost, proportional to the gene copy number, mRNA copy number, and protein copy number of the exogenous gene, respectively. The arrow widths approximately represent their relative contributions to the total fitness cost (Table I).

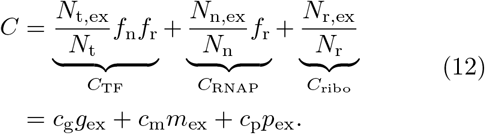

The transcription cost now consists of two components generated by competition for TF and RNAP: *C*_tx_ = *C*_TF_+*C*_RNAP_ (Figure 3b). Here, *N*_t,ex_ is the copy number of TF working on the exogenous gene, and *N*_t_ is the total copy number of TF. One should note that the TF cost is damped by the two downstream factors *f*_n_ and *f*_r_. We introduce the TF cost per gene as *c*_*g*_ = Λ_t,ex_*P*_t,ex_*f*_n_*f*_r_*/N*_t_ where Λ_t,ex_ is the maximum TF number recruited by one copy of the exogenous gene and *P*_t,ex_ is the corresponding TF binding probability. The TF cost shows that introducing an exogenous gene into the genome alone already imposes a cost due to TF competition, even without downstream mRNA production.

Because it is difficult to estimate *N*_t,ex_ and *N*_t_ directly, we take an indirect approach assuming that the TF binding limits the mRNA production rates, that is, the ratio between the mRNA production rates of the exogenous gene and other genes can be approximated by the corresponding ratio of the copy numbers of TFs working on each promoter, *β*_m,ex_*/β*_m_ ≈ (*N*_t,ex_*/g*_ex_)*/*(*N*_t_(1 − *f*_t_)*/g*) where *f*_t_ is the fraction of free TFs. Given this approximation, we estimate *c*_g_ in Eq. (12) as

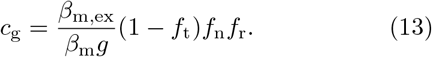

Intriguingly, the estimated TF cost is close to the measured transcription cost (Table I), meaning that the TF cost dominates the total transcription cost. To summarize, our theory decomposes the total fitness cost into different components. In particular, the translation cost is dominated by competition for ribosomes, and the transcription cost is dominated by competition for TF (Figure 3b).

### Effects of gene constructs and growth conditions

We rewrite Eq. (12) as

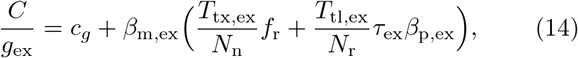

where *β*_m,ex_ is the mRNA production rate per copy of the exogenous gene and *β*_p,ex_ is the protein production rate per copy of the exogenous mRNA. Eq. (14) predicts that the cost per gene copy positively correlates with the mRNA production rate *β*_m,ex_. We compare our predictions with experimental data for *S. cerevisiae* [8], where the control is the wild-type mCherry gene driven by the TDH3 promoter in standard culture (SC). In agreement with our predictions, the construct with the promoter PGK1, which is weaker than TDH3, has a lower cost in SC (Figure 4a). Eq. (14) also predicts that if the produced mRNA has a shorter lifetime *τ*_ex_, the cost per gene copy should also be smaller, and this prediction agrees with the lower cost of the DAmP mutant of mCherry mRNA [8], whose lifetime is about 1/10 of the control (Figure 4a). Since the protein degradation rate does not enter Eq. (14), we also predict that the cost per gene copy is independent of protein stability, as validated by the experiments showing that the protein variant (CLN2) with a much higher degradation rate has a cost per gene copy similar to that of the control (Figure 4a).

**FIG. 4.**
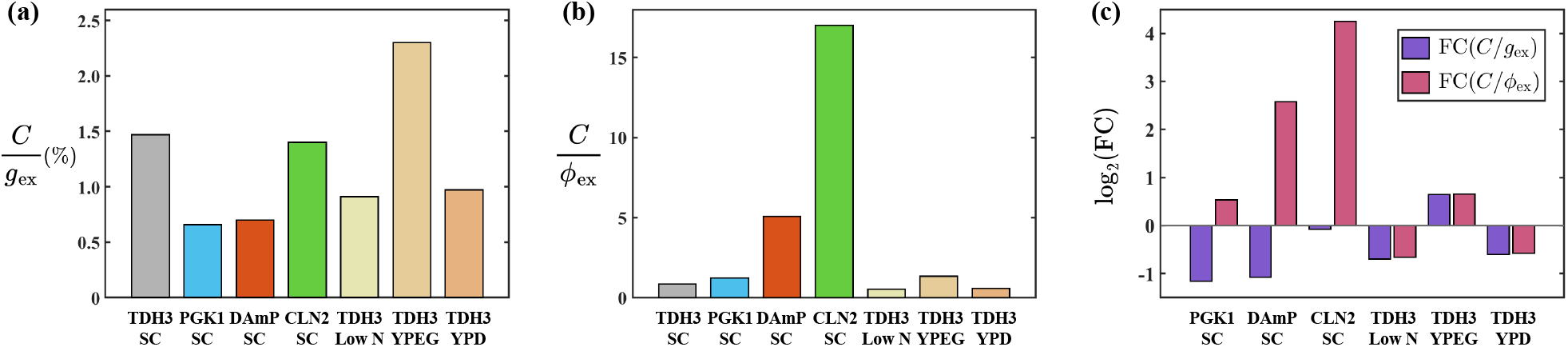
The fitness cost of *S. cerevisiae* for different gene constructs and growth conditions. (a) The experimentally measured fitness cost per gene copy number (*g*_ex_). We use the wild-type mCherry gene under the TDH3 promoter in standard culture (SC) as the control. PGK1 is a weaker promoter compared to TDH3. The DAmP construct produces an unstable mRNA with a shorter lifetime. The CLN2 construct produces a protein with a high degradation rate (note that in the experiments of Ref. [8], the CLN2 construct in fact produced GFP, which is practically identical to mCherry regarding fitness cost). Low N represents a nitrogen-limited medium, where the amount of transcriptional resources is measured to be higher than SC [31]. YPEG medium lacks fermentable carbon sources, while YPD is a rich medium with a complete nutrient supply. (b) The experimentally measured fitness cost per proteome fraction. (c) The fold change (FC) of the cost per gene copy and per proteome fraction compared to the control (TDH3, SC).

We also calculate the cost per proteome fraction generated by the exogenous gene (Supplementary Information Section B), a metric often used in experiments for both *S. cerevisiae* and *E. coli* (Table S3):

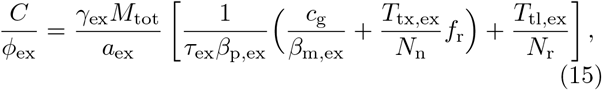

where *M*_tot_ is the total protein mass, *ϕ*_ex_ is the proteome fraction of the exogenous proteins. Interestingly, our model predicts that the PGK1 construct with a lower *β*_m,ex_, the DAmP construct with a shorter *τ*_ex_, and the CLN2 construct with a higher protein degradation rate *γ*_ex_ than TDH3, should all have a higher cost per proteome fraction than TDH3. All these predictions agree with experiments (Figure 4b).

When altering growth conditions, the cost per gene copy and the cost per proteome fraction change differently. In the low N condition where cells are richer in transcriptional resources [31], we expect a smaller *c*_g_ (the TF cost per gene copy, Eq. (12)) and a larger *N*_n_ (the total number of RNAP), corresponding to a lower cost both per gene copy and per proteome fraction. This is in agreement with the experimental data for the control and the same construct in the low N condition (Figure 4a-b). For the nutrient-poor medium YPEG, the transcription and translation elongation speeds are presumably slower, leading to a longer *T*_tx,ex_ and *T*_tl,ex_. Therefore, we predict that the cost per gene copy and per proteome fraction should both be higher for TDH3 in YPEG than the control, i.e., TDH3 in SC. In contrast, for the rich medium YPD, we expect a shorter *T*_tx,ex_ and *T*_tl,ex_, and the cost per gene copy and per proteome fraction should both be lower for TDH3 in YPD than SC (Figure 4a-b).

We also calculate the fold change (FC) of different gene constructs and conditions compared to the control, which clearly illustrates that the cost per gene copy and the cost per proteome fraction move in opposite directions for different gene constructs but move in the same direction when the medium changes (Figure 4c).

### Cost generated by the products is negligible

So far, we have considered a fixed resource pool with constant resource copy numbers and cell volume, which can be a good approximation when the exogenous protein is quickly degraded. For non-degradable proteins, the cell volume can increase upon expression of an exogenous gene [4, 8, 48], which may dilute key factors such as ribosomes and RNAP. On the other hand, cells also make more endogenous proteins to compensate for the dilution effects [8]. We extend our model to a more general scenario where the resource copy numbers (i.e., RNAP number *N*_n_ and ribosome number *N*_r_) and cell volume change due to the expression of the exogenous gene, which may generate an additional fitness cost. We refer to this additional fitness cost as the cost of the products (*C*^*′*^, as opposed to the cost in the processes of transcription and translation), which is found to be (Supplementary Information Section C):

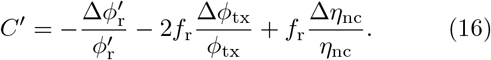

Here, 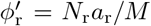, the ribosome proteome fraction in the endogenous proteins, different from *ϕ*_r_ = *N*_r_*a*_r_*/M*_tot_, the ribosome proteome fraction in the total proteome. *ϕ*_tx_ is the proteome fraction encoding transcriptional proteins (e.g., RNAP and transcription factors). *η*_nc_ = *V*_n_*/V*_c_ is the volume ratio of nucleus to cytoplasm (N/C ratio). 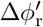, Δ*ϕ*_tx_, and Δ*η*_nc_ in Eq. (16) represent the active/passive regulation of translation and transcription resources, and the N/C ratio. Notably, Eq. (16) illustrates that increasing translation and transcription resources can relieve the cost, but the rise of the nucleus-to-cytoplasm volume ratio aggravates the cost.

Intriguingly, experimental data from *S. cerevisiae* show that although *ϕ*_r_ is diluted by the exogenous proteins produced, 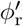 stays almost invariant (Figure 5). Furthermore, the proteome fraction encoding transcriptional proteins is also approximately constant [31], which implies that the volume expansion is compensated for by making more transcriptional proteins. We expect that the N/C ratio is also approximately constant if the protein products of the exogenous gene are evenly distributed in the cell [49]. Therefore, according to Eq. (16), the fitness cost generated by the product is insignificant compared to the overall fitness cost, consistent with previous experimental observations in *E. coli* [12] and *S. cerevisiae* [8]. Here, we point out that this negligible cost of the products applies to unnecessary proteins without any specific function or toxicity when overexpressed. For proteins that can change the nutrient uptake (e.g., LacY [50]) or are toxic due to misfolding [51–53], the products naturally generate a considerable fitness cost, which should be isolated as a protein-specific cost.

**FIG. 5.**
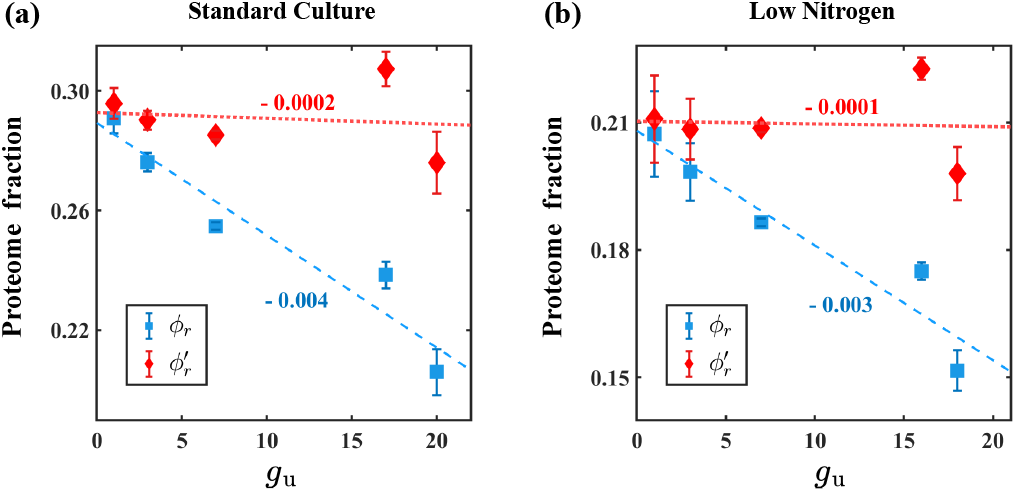
The ribosomal proteome fraction in the total proteome (*ϕ*_r_) and the ribosomal proteome fraction in the endogenous proteome 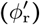. With the expression of mCherry proteins in *S. cerevisiae, ϕ*_r_ decreases, whereas 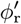 stays almost constant. The dashed lines show the linear fits to the data points, with the numbers indicating the slopes. Data are taken from Ref. [31], and the results under the two experimental conditions show consistency.

## DISCUSSION

Many experiments have measured the growth rate reduction to quantify the fitness cost of gene expression [1, 2, 4, 8]. It has been widely presumed that the cost arises from both transcription and translation [8, 10, 13]. However, the mechanism behind this hypothesis remains poorly understood. In this work, we use a simple resource competition model and disentangle the fitness cost into different components generated by TF, RNAP, and ribosome competition. In particular, we show that the ribosome cost dominates the translation cost, and the transcription cost in *S. cerevisiae* is dominated by the TF cost. We remark that the coarse-grained TF in our model may correspond to the general transcription factors, which help to position RNAP correctly at the promoters, aid in separating the two strands of DNA, and release RNAP from the promoters to start elongation, including TFIIB, TFIID, etc. These proteins are needed at nearly all promoters used by RNA polymerase II [46, 47]. Other crucial proteins may also be candidates for the limiting factors, including mediators [45], transcription activators, and chromatin remodeling proteins [47]. Our framework provides valuable guidance on designing gene circuits to minimize interference with the host cells [54, 55]. We also discuss the differences between our model and previous ones regarding the fitness cost [4, 10, 11, 13] in Supplementary Information Section A and Table S1-S2.

We propose more systematic experiments in the future to obtain a deeper understanding of the fitness cost of gene expression. First, measurements of the simultaneous changes in the cell volume, ribosome copy number, RNAP copy number, etc., upon gene overexpression will be invaluable for us to understand how cells adapt to limited resources for gene expression. Second, more accurate and high-throughput measurements of fitness cost are required to enable quantitative modeling with more mechanistic insights.

## METHODS

### A. Derivation of the fitness cost

The probability of an endogenous promoter bound by an RNAP is

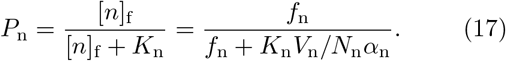

Here, [*n*]_f_ = *N*_n_*f*_n_*α*_n_*/V*_n_ is the concentration of free RNAP in the nucleus; *α*_n_ is the fraction of free RNAP in the inactive RNAP pool, which we set as a constant, as it is determined by molecular binding affinities (e.g., non-specific binding affinity); *V*_n_ is the nuclear volume; and *K*_n_ is the corresponding dissociation constant of the endogenous genes [16–18]. Differentiating Eq. (17) leads to

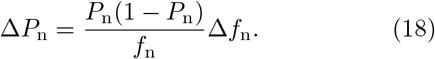

Combining the above equation with Eq. (2), we find that

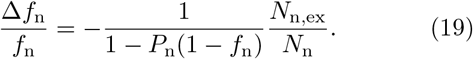

In deriving Eq. (19), we have used the approximation *N*_n,ex_ ≪ *N*_n_, which holds in biological scenarios where a single gene copy occupies a small fraction of total RNAPs.

The probability for the ribosome-binding-site of endogenous mRNA to be bound by a ribosome is

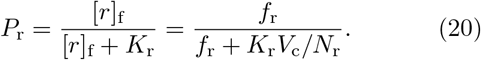

Here, [*r*]_f_ = *N*_r_*f*_r_*/V*_c_ is the concentration of free ribosomes in the cytoplasm, *V*_c_ is the cytoplasmic volume, and *K*_r_ is the corresponding dissociation constant of endogenous mRNAs [14, 15]. Similarly, differentiating Eq. (20) and combining it with Eqs. (2, 3, 5) lead to

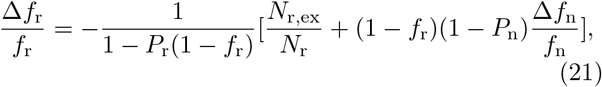

where *N*_r,ex_ is the number of ribosomes on the exogenous mRNA.

Differentiating Eq. (7), we get

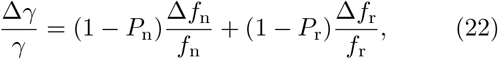

where *γ* = *µ* + *d* is the protein decay rate of the average endogenous protein, the sum of the growth rate and the average protein degradation rate. Given the definition of cost (Eq. (8)), and combining Eqs. (19) and (21-22), we get

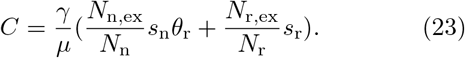

Here, we introduce *s*_n_ and *s*_r_ as the sensitivity factors

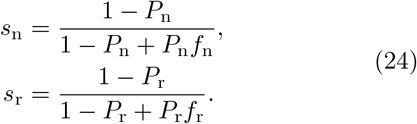

In the limit *P*_n_ →0, transcription activity is rare, and *s*_n_ reaches its maximum value 1, meaning that transcription initiation is sensitive to changes in the RNAP resources.

On the contrary, in the limit *P*_n_ → 1, cells have plenty of RNAPs and *s*_n_ = 0, meaning that transcription is RNAP-saturated. A similar discussion applies to *s*_r_. We also introduce the downstream factor

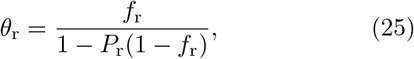

which represents the dampening effects of downstream processes on the RNAP cost: a small fraction of free ribosomes *f*_r_ dampens the RNAP cost.

In typical biological scenarios applied to microorganisms, Eq. (23) is simplified to Eq. (9) under the following conditions: 1. *γ* ≈*µ* because endogenous protein degradation is negligible compared to the growth rate [25–27]; 2. *P*_n_ is significantly lower than one such that *s*_n_ ≈ 1; 3. *P*_r_ is significantly lower than one such that *s*_r_ ≈ 1 and *θ*_r_ ≈ *f*_r_. We remark that the conditions *P*_n_ ≪1 and *P*_r_ ≪1 are only the properties of the “average” endogenous gene, which is typically weakly expressed [13, 18, 20, 56].

At steady state, the mRNA degradation rate equals the number of transcription initiation events per unit time, leading to

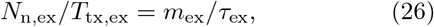

where *τ*_ex_ is the exogenous mRNA lifetime, and *T*_tx,ex_ is the time for an RNAP to transcribe the exogenous gene. Similarly, from the equilibrium between the number of proteins diluted and degraded per unit time and the translation initiation events per unit time, we have

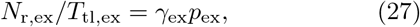

where *γ*_ex_ = *µ* + *d*_ex_ is the protein decay rate of the exogenous gene. Replacing the *N*_n,ex_ and *N*_r,ex_ in Eq. (9) by Eqs. (26-27), we finally get Eq. (10).

### B. Estimating the transcription and translation cost per gene/mRNA/protein from experiments in *S. cerevisiae*

The fitness costs of two gene constructs expressing mCherry proteins were measured, denoted as “wt” and “DAmP”, respectively [8]. Both constructs were driven by the TDH3 promoter. However, DAmP lacks a terminator, which leads to a shorter mRNA lifetime (*τ*_DAmP_) than the wt mRNA lifetime (*τ*_wt_), and further leads to a lower copy number of mRNA and a lower copy number of protein. Assuming all other parameters are unaffected, we have *m*_DAmP_*/m*_wt_ = *p*_DAmP_*/p*_wt_ = *τ*_DAmP_*/τ*_wt_. Both Eq. (10) and Eq. (14) show that the transcription costs of the two constructs are the same, but the translation costs differ by a factor of *τ*_DAmP_*/τ*_wt_. Therefore, we have the following linear equations:

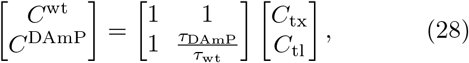

where *C*^wt^ and *C*^DAmP^ are experimentally measured fitness cost per gene copy of the wt and the DAmP construct, respectively. Here, *C*_tx_ and *C*_tl_ refer to the transcription and translation costs of the wt construct, respectively. Solving the simultaneous linear equations with *C*^wt^*/g*_ex_ = 1.50 ×10^*−*2^, *C*^DAmP^*/g*_ex_ = 7.0 ×10^*−*3^, and *τ*_DAmP_*/τ*_wt_ = 0.1, we get *C*_tl_*/g*_ex_ = 8.9× 10^*−*3^ and *C*_tx_*/g*_ex_ = 6.1 ×10^*−*3^. Here, *C*^wt^*/g*_ex_ and *C*^DAmP^*/g*_ex_ are obtained from the linear fitting of the experimental data [8].

Each gene copy contributes about 1.7% to the proteome for the TDH3 construct in the standard culture [8], which corresponds to a protein copy number of *p*_ex_*/g*_ex_ = 1.7% × *M/a*_ex_ = 1.5 ×10^6^ with *M* = 4.0 ×10^*−*12^ g [57, 58] and *a*_ex_ = 27 kDa [59]. Assuming that the protein production rate per mRNA of the exogenous gene is the same as that of the average endogenous genes (*β*_p,ex_ = 1 ×10^3^/h [60]), we estimate *m*_ex_*/g*_ex_ = *γ*_ex_*p*_ex_*/β*_p,ex_*g*_ex_ = 6 ×10^2^ with *γ*_ex_ = 0.4/h [8].

### C. The fitness cost due to transcription factors

To include the effects of TFs, we write the conservation equation for transcription factors in the absence of the exogenous gene:

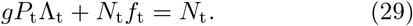

Here, Λ_t_ is the maximum number of TF on an endogenous promoter, and *P*_t_ is the probability for an endogenous promoter to be bound by TF, which is TFconcentration dependent: 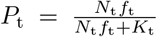, with *K*_t_ the coarse-grained dissociation constant. The copy number of RNAPs on each gene is Λ_n_*P*_t_*P*_n_, and the endogenous mRNA production rate is 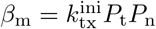. Similar results apply to the exogenous gene. By including the exogenous gene, and following the same method of derivation as in Methods A, we find the fitness cost as

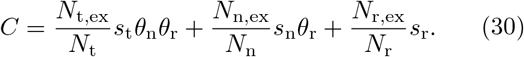

Here, *N*_t,ex_ is the number of TF on the promoter for the exogenous gene. The sensitivity factor of TF is

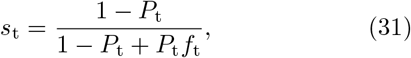

and the downstream factor is

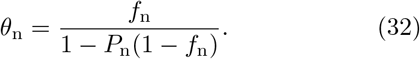

Notably, the total fitness cost can be precisely decomposed into the contributions of each limiting resource. The corresponding downstream resources, including RNAP and ribosomes, dampen the cost of a finite number of TF. Approximating *s*_t_ = *s*_n_ = *s*_r_ = 1, *θ*_n_ = *f*_n_ and *θ*_r_ = *f*_r_ under *P*_t_, *P*_n_ ≪ 1, we obtain Eq. (12).

## ACKNOWLEDGMENTS

The research was funded by the National Key Research and Development Program of China (2024YFA0919600) and supported by Peking-Tsinghua Center for Life Sciences grants.

## References

[1] W. E. Bentley, N. Mirjalili, D. C. Andersen, R. H. Davis, and D. S. Kompala, Plasmid-encoded protein: the principal factor in the metabolic burden associated with recombinant bacteria, Biotechnology and bioengineering 35, 668 (1990).

[2] H. Dong, L. Nilsson, and C. G. Kurland, Gratuitous over-expression of genes in escherichia coli leads to growth inhibition and ribosome destruction, Journal of Bacteriology 177, 1497 (1995).

[3] C. Rang, J. E. Galen, J. B. Kaper, and L. Chao, Fitness cost of the green fluorescent protein in gastrointestinal bacteria, Canadian journal of microbiology 49, 531 (2003).

[4] M. Scott, C. W. Gunderson, E. M. Mateescu, Z. Zhang, and T. Hwa, Interdependence of cell growth and gene expression: origins and consequences, Science 330, 1099 (2010).

[5] M. Lynch and G. K. Marinov, The bioenergetic costs of a gene, Proceedings of the National Academy of Sciences 112, 15690 (2015).

[6] A. San Millan, M. Toll-Riera, Q. Qi, and R. C. MacLean, Interactions between horizontally acquired genes create a fitness cost in pseudomonas aeruginosa, Nature communications 6, 6845 (2015).

[7] Y. Eguchi, K. Makanae, T. Hasunuma, Y. Ishibashi, K. Kito, and H. Moriya, Estimating the protein burden limit of yeast cells by measuring the expression limits of glycolytic proteins, eLife 7, e34595 (2018).

[8] M. Kafri, E. Metzl-Raz, G. Jona, and N. Barkai, The cost of protein production, Cell reports 14, 22 (2016).

[9] A. Wagner, Energy constraints on the evolution of gene expression, Molecular Biology and Evolution 22, 1365 (2005).

[10] T. W. Lo, H. J. Choi, D. Huang, and P. A. Wiggins, Noise robustness and metabolic load determine the principles of central dogma regulation, Science Advances 10, eado3095 (2024).

[11] L. Calabrese, L. Ciandrini, and M. Cosentino Lagomarsino, How total mrna influences cell growth, Proceedings of the National Academy of Sciences 121, e2400679121 (2024).

[12] D. M. Stoebel, A. M. Dean, and D. E. Dykhuizen, The cost of expression of escherichia coli lac operon proteins is in the process, not in the products, Genetics 178, 1653 (2008).

[13] J. Hausser, A. Mayo, L. Keren, and U. Alon, Central dogma rates and the trade-off between precision and economy in gene expression, Nature communications 10, 68 (2019).

[14] J. Lin and A. Amir, Homeostasis of protein and mrna concentrations in growing cells, Nature communications 9, 4496 (2018).

[15] Q. Wang and J. Lin, Heterogeneous recruitment abilities to rna polymerases generate nonlinear scaling of gene expression with cell volume, Nature communications 12, 6852 (2021).

[16] S. Klumpp and T. Hwa, Growth-rate-dependent partitioning of rna polymerases in bacteria, Proceedings of the National Academy of Sciences 105, 20245 (2008).

[17] M. P. Swaffer, G. K. Marinov, H. Zheng, L. F. Valenzuela, C. Y. Tsui, A. W. Jones, J. Greenwood, A. Kundaje, W. J. Greenleaf, R. Reyes-Lamothe, et al., Rna polymerase ii dynamics and mrna stability feedback scale mrna amounts with cell size, Cell 186, 5254 (2023).

[18] Y. Yan, T. Li, and J. Lin, Buffering effects of nonspecifically dna-bound rna polymerases in bacteria, Physical Review Research 6, 033133 (2024).

[19] P. Shah, Y. Ding, M. Niemczyk, G. Kudla, and J. B. Plotkin, Rate-limiting steps in yeast protein translation, Cell 153, 1589 (2013).

[20] O. Borkowski, F. Ceroni, G.-B. Stan, and T. Ellis, Overloaded and stressed: whole-cell considerations for bacterial synthetic biology, Current opinion in microbiology 33, 123 (2016).

[21] F. Ceroni, R. Algar, G.-B. Stan, and T. Ellis, Quantifying cellular capacity identifies gene expression designs with reduced burden, Nature methods 12, 415 (2015).

[22] I. Frumkin, D. Schirman, A. Rotman, F. Li, L. Zahavi, E. Mordret, O. Asraf, S. Wu, S. F. Levy, and Y. Pilpel, Gene architectures that minimize cost of gene expression, Molecular Cell 65, 142 (2017).

[23] U. Raue, S. Oellerer, and S. Rospert, Association of protein biogenesis factors at the yeast ribosomal tunnel exit is affected by the translational status and nascent polypeptide sequence, Journal of Biological Chemistry 282, 7809 (2007).

[24] E. M. Phizicky and A. K. Hopper, trna biology charges to the front, Genes & development 24, 1832 (2010).

[25] R. Christiano, N. Nagaraj, F. Fröhlich, and T. C. Walther, Global proteome turnover analyses of the yeasts s. cerevisiae and s. pombe, Cell reports 9, 1959 (2014).

[26] M. Martin-Perez and J. Villén, Determinants and regulation of protein turnover in yeast, Cell Systems 5, 283 (2017).

[27] R. Christiano, H. Arlt, S. Kabatnik, N. Mejhert, Z. W. Lai, R. V. Farese, and T. C. Walther, A systematic protein turnover map for decoding protein degradation, Cell reports 33 (2020).

[28] W. Qian, J.-R. Yang, N. M. Pearson, C. Maclean, and J. Zhang, Balanced codon usage optimizes eukaryotic translational efficiency, PLoS genetics 8, e1002603 (2012).

[29] A. Riba, N. Di Nanni, N. Mittal, E. Arhńe, A. Schmidt, and M. Zavolan, Protein synthesis rates and ribosome occupancies reveal determinants of translation elongation rates, Proceedings of the National Academy of Sciences 116, 15023 (2019).

[30] T. von der Haar, A quantitative estimation of the global translational activity in logarithmically growing yeast cells, BMC systems biology 2, 87 (2008).

[31] E. Metzl-Raz, M. Kafri, G. Yaakov, I. Soifer, Y. Gurvich, and N. Barkai, Principles of cellular resource allocation revealed by condition-dependent proteome profiling, eLife 6, e28034 (2017).

[32] T. Borggrefe, R. Davis, A. Bareket-Samish, and R. D. Kornberg, Quantitation of the rna polymerase ii transcription machinery in yeast, Journal of Biological Chemistry 276, 47150 (2001).

[33] C. Miller, B. Schwalb, K. Maier, D. Schulz, S. Dümcke, B. Zacher, A. Mayer, J. Sydow, L. Marcinowski, L. Dölken, et al., Dynamic transcriptome analysis measures rates of mrna synthesis and decay in yeast, Molecular systems biology 7, 458 (2011).

[34] B. Neymotin, R. Athanasiadou, and D. Gresham, Determination of in vivo rna kinetics using rate-seq, Rna 20, 1645 (2014).

[35] P. Eser, C. Demel, K. C. Maier, B. Schwalb, N. Pirkl, D. E. Martin, P. Cramer, and A. Tresch, Periodic mrna synthesis and degradation co-operate during cell cycle gene expression, Molecular systems biology 10, 717 (2014).

[36] A. M. Edwards, C. M. Kane, R. A. Young, and R. D. Kornberg, Two dissociable subunits of yeast rna polymerase ii stimulate the initiation of transcription at a promoter in vitro, Journal of Biological Chemistry 266, 71 (1991).

[37] T. O’Brien and J. T. Lis, Rapid changes in drosophila transcription after an instantaneous heat shock, Molecular and cellular biology 13, 3456 (1993).

[38] J.E. Pérez-Ortín, P. M. Alepuz, and J. Moreno, Genomics and gene transcription kinetics in yeast, TrendS in Genetics 23, 250 (2007).

[39] A. Goffeau, B. G. Barrell, H. Bussey, R. W. Davis, B. Dujon, H. Feldmann, F. Galibert, J. D. Hoheisel, C. Jacq, M. Johnston, et al., Life with 6000 genes, Science 274, 546 (1996).

[40] K. P. Byrne and K. H. Wolfe, The yeast gene order browser: combining curated homology and syntenic context reveals gene fate in polyploid species, Genome research 15, 1456 (2005).

[41] V. Pelechano, S. Chavez, and J. E. Perez-Ortin, A complete set of nascent transcription rates for yeast genes, PloS one 5, e15442 (2010).

[42] P. Shah and M. Gilchrist, Explaining complex codon usage patterns with selection for translational efficiency, mutation bias, and genetic drift, Proceedings of the National Academy of Sciences of the United States of America 108, 10231 (2011).

[43] Mahima and A. K. Sharma, Optimization of ribosome utilization in saccharomyces cerevisiae, PNAS Nexus 2, pgad074 (2023).

[44] E. Metzl-Raz, M. Kafri, G. Yaakov, and N. Barkai, Gene transcription as a limiting factor in protein production and cell growth, G3: Genes, Genomes, Genetics 10, 3229 (2020).

[45] Y. Takagi and R. D. Kornberg, Mediator as a general transcription factor, Journal of Biological Chemistry 281, 80 (2006).

[46] S. Hahn and E. T. Young, Transcriptional regulation in saccharomyces cerevisiae: transcription factor regulation and function, mechanisms of initiation, and roles of activators and coactivators, Genetics 189, 705 (2011).

[47] B. Alberts, R. Heald, A. Johnson, D. Morgan, M. Raff, K. Roberts, and P. Walter, Molecular biology of the cell (WW Norton & Company, 2022).

[48] M. Basan, M. Zhu, X. Dai, M. Warren, D. Sévin, Y.P. Wang, and T. Hwa, Inflating bacterial cells by increased protein synthesis, Molecular systems biology 11, 836 (2015).

[49] J. Lemiére, P. Real-Calderon, L. J. Holt, T. G. Fai, and F. Chang, Control of nuclear size by osmotic forces in schizosaccharomyces pombe, eLife 11, e76075 (2022).

[50] M. Eames and T. Kortemme, Cost-benefit tradeoffs in engineered lac operons, Science 336, 911 (2012).

[51] D. Allan Drummond and C. O. Wilke, The evolutionary consequences of erroneous protein synthesis, Nature Reviews Genetics 10, 715 (2009).

[52] K. A. Geiler-Samerotte, M. F. Dion, B. A. Budnik, S. M. Wang, D. L. Hartl, and D. A. Drummond, Misfolded proteins impose a dosage-dependent fitness cost and trigger a cytosolic unfolded protein response in yeast, Proceedings of the National Academy of Sciences 108, 680 (2011).

[53] Z. Farkas, D. Kalapis, Z. Bodi, B. Szamecz, A. Daraba, K. Almasi, K. Kovacs, G. Boross, F. Pal, P. Horvath, et al., Hsp70-associated chaperones have a critical role in buffering protein production costs, eLife 7, e29845 (2018).

[54] O. Borkowski, A. Goelzer, M. Schaffer, M. Calabre, U. Mäder, S. Aymerich, M. Jules, and V. Fromion, Translation elicits a growth rate-dependent, genome-wide, differential protein production in bacillus subtilis, Molecular systems biology 12, 870 (2016).

[55] A. Grob, R. Di Blasi, and F. Ceroni, Experimental tools to reduce the burden of bacterial synthetic biology, Current Opinion in Systems Biology 28, 100393 (2021).

[56] R. Balakrishnan, M. Mori, I. Segota, Z. Zhang, R. Aebersold, C. Ludwig, and T. Hwa, Principles of gene regulation quantitatively connect dna to rna and proteins in bacteria, Science 378, eabk2066 (2022).

[57] G. C. Johnston, J. R. Pringle, and L. H. Hartwell, Coordination of growth with cell division in the yeast saccharomyces cerevisiae, Experimental cell research 105, 79 (1977).

[58] B. Futcher, G. Latter, P. Monardo, C. McLaughlin, and J. Garrels, A sampling of the yeast proteome, Molecular and cellular biology 19, 7357 (1999).

[59] M. Sette, L. A. Johnson, R. Jimenez, and F. A. Mulder, Backbone 1H, 15N and 13C resonance assignments of the 27kDa fluorescent protein mcherry, Biomolecular NMR Assignments 17, 243 (2023).

[60] D. E. Weinberg, P. Shah, S. W. Eichhorn, J. A. Hussmann, J. B. Plotkin, and D. P. Bartel, Improved ribosome-footprint and mrna measurements provide insights into dynamics and regulation of yeast translation, Cell reports 14, 1787 (2016).

