## Supplementary Information for "Disentangling the fitness cost of gene expression"

### A. Comparison between our model and others

In Ref. [1], Hausser et al. supposed that the translation cost of expressing an extra gene is $C_{\rm tl} = p_{\rm ex}/p$, and the transcription cost is $C_{\rm tx} = m_{\rm ex}/m$, where $p_{\rm ex}$ and $m_{\rm ex}$ are the copy numbers of exogenous proteins and mRNAs; $p$ and $m$ are the copy numbers of endogenous proteins and mRNAs.

In Ref. [2], Lo et al. proposed a concept called the effective total metabolic load: $L_{\rm eff} = p + \lambda m'$, where $\lambda$ is a factor calibrating the load per mRNA produced to the load per protein produced. $m' = \frac{T}{\tau} m$ is the endogenous mRNA number produced per cell cycle. The authors assumed that the proteins are non-degradable; therefore, the endogenous protein number $p$ is also the protein number produced per cell cycle. The translation cost of the exogenous gene is $C_{\rm tl} = \frac{p_{\rm ex}}{L_{\rm eff}}$, and the transcription cost is $C_{\rm tx} = \frac{\lambda m'_{\rm ex}}{L_{\rm eff}}$, where $m'_{\rm ex} = \frac{T}{\tau_{\rm ex}} m_{\rm ex}$ is the exogenous mRNA number produced per cell cycle.

In Ref. [3], Scott et al. assumed that the growth rate $\mu$ is proportional to $1 - \phi_{\rm Q} - \phi_{\rm r,0} - \phi_{\rm ex}$, where $\phi_{\rm Q}$ is a constant proteome fraction of the housekeeping sector Q, $\phi_{\rm r,0}$ is a constant proteome fraction of ribosomal proteins that do not contribute to growth, and $\phi_{\rm ex}$ is the proteome fraction of the extra protein. The fitness cost comes from the dilution of ribosome fraction due to the expression of the exogenous genes, which is $C = -\frac{\Delta\mu}{\mu} = \frac{\phi_{\rm ex}}{1-\phi_{\rm Q}-\phi_{\rm r,0}}$.

In Ref. [4], Calabrese et al. assumed that the growth rate $\mu$ is proportional to $\phi_{\rm r} f_{\rm b,r}$, where $f_{\rm b,r} = 1 - f_{\rm r} = \frac{\phi_{\rm n} f_{\rm b,n}}{\phi_{\rm n} f_{\rm b,n} + K_{\rm eff}}$ is the fraction of bound ribosomes, with $f_{\rm b,n} = 1 - f_{\rm n}$ the fraction of bound RNAPs. The expression of extra genes leads to a proteome fraction $\phi_{\rm ex}$, which influences $\mu$ by changing $\phi_{\rm r}$, $\phi_{\rm n}$, and $f_{\rm b,n}$. The fitness cost $C = -\frac{\Delta\mu}{\mu} = -[\frac{\Delta\phi_{\rm r}}{\phi_{\rm r}} + f_{\rm r}(\frac{\Delta\phi_{\rm n}}{\phi_{\rm n}} + \frac{\Delta f_{\rm b,n}}{f_{\rm b,n}})]$. Assuming a constant proteome fraction ($\phi_{\rm Q}$) of the housekeeping sector Q, they have $\frac{\Delta\phi_{\rm r}}{\phi_{\rm r}} = -\frac{\phi_{\rm ex}}{1-\phi_{\rm Q}}$ due to dilution effects; $\frac{\Delta\phi_{\rm n}}{\phi_{\rm n}} = -\frac{\phi_{\rm ex}}{1-\phi_{\rm Q}}$ due to dilution effects if $n \notin {\rm Q}$ or $\frac{\Delta\phi_{\rm n}}{\phi_{\rm n}} = 0$ if $n \in {\rm Q}$; $\frac{\Delta f_{\rm b,n}}{f_{\rm b,n}} = f_{\rm n}\frac{\phi_{\rm ex}}{1-\phi_{\rm Q}}$ because the insertion of extra genes increases the fraction of bound RNAPs (see the details of derivation in the supporting information of [4]). According to the above information, we have $\frac{C}{\phi_{\rm ex}} = \frac{1}{1-\phi_{\rm Q}}[1+f_{\rm r}(1-f_{\rm n})]$ ($n \notin {\rm Q}$) or $\frac{C}{\phi_{\rm ex}} = \frac{1}{1-\phi_{\rm Q}}[1 - f_{\rm r} f_{\rm n}]$ ($n \in {\rm Q}$).

We remark that in [1] and [2], the fitness costs essentially come from the processes of gene expression where various resources are consumed; in [3] and [4], the fitness costs primarily arise from the dilution effects of expressed protein products. Our model considers the costs both in the processes and the products, and finds that the transcriptional and translational processes primarily account for the fitness cost. A summary of the above four models is presented in Table S1. A quantitative comparison between models and experiments is presented in Table S2. Further experiments are needed to systematically validate these models.

TABLE S1. **A summary of the four other models on the fitness cost of gene expression.**

| | **Hausser et al. [1]** | **Lo et al. [2]** | **Scott et al. [3]** | **Calabrese et al. [4]** |
|---|---|---|---|---|
| Primary origin of cost | Processes | Processes | Products | Products |
| $C$ | $\frac{p_{\rm ex}}{p}+\frac{m_{\rm ex}}{m}$ | $\frac{p_{\rm ex}+\lambda m'_{\rm ex}}{p+\lambda m'}$ | $\frac{\phi_{\rm ex}}{1-\phi_{\rm Q}-\phi_{\rm r,0}}$ | $\frac{1+f_{\rm r}(1-f_{\rm n})}{1-\phi_{\rm Q}}\phi_{\rm ex}$ |
| $\frac{C_{\rm tl}}{p_{\rm ex}}$ | $\frac{1}{p}$ | $\frac{1}{p+\lambda m'}$ | —— | —— |
| $\frac{C_{\rm tx}}{m_{\rm ex}}$ | $\frac{1}{m}$ | $\frac{\lambda T}{\tau_{\rm ex}}\frac{1}{p+\lambda m'}$ | —— | —— |

TABLE S2. **A quantitative comparison among theories and experimental results.** Here, we analyze the data of the mCherry gene driven by TDH3 promoter in the standard culture. Details on the calculations of a-b are explained in Table I and Methods B in the main text. Values of c-f are calculated according to expressions in Table S1. The parameter values we use are as follows (if present): $a_{\rm ex}=27$ kDa [5], $a=50$ kDa [6, 7], $M=4.0\times10^{-12}$ g [8, 9], $\phi_{\rm ex}/g_{\rm ex}=1.7\%$, $p_{\rm ex}/g_{\rm ex}=1.5\times10^{6}$, $m_{\rm ex}/g_{\rm ex}=6\times10^{2}$ (Methods B in the main text), $p=M/a=5\times10^{7}$, $m=3\times10^{4}$ [10, 11], $T=1.7$ h [12], $\tau=\tau_{\rm ex}=15$ min [13–15], $\phi_{\rm Q}=0.2$ [4], $\phi_{\rm r,0}=0.08$ [16], $f_{\rm r}=0.3$ [16], $f_{\rm n}=0.9$ [11]. The corresponding expressions of (c) and (d) are multiplied by $a_{\rm ex}/a=0.54$, taking account of the shorter mRNA and protein length of a mCherry gene compared to an average endogenous gene. The calibration factor $\lambda$ is set to be 100 or 10 in (d) [2], corresponding to two results separated by a comma. The two values in (f) represent the cases $n\notin$ Q and $n\in$ Q, respectively. Here, all values are kept to one decimal place for the convenience of comparison. A dash means that the corresponding value is not included in the theory or experiment.

| **Value** | $\frac{C}{\phi_{\rm ex}}$ | $\frac{C_{\rm tl}}{p_{\rm ex}}$ $(10^{-8})$ | $\frac{C_{\rm tx}}{m_{\rm ex}}$ $(10^{-5})$ | $\frac{C_{\rm RNAP}}{m_{\rm ex}}$ $(10^{-7})$ | $\frac{C_{\rm TF}}{g_{\rm ex}}$ $(10^{-3})$ |
|---|---|---|---|---|---|
| **(a) Experiment** | 0.9 | 0.6 | 1.0 | —— | —— |
| **(b) This article** | 1.1 | 0.9 | 0.8 | 3.7 | 4.5 |
| **(c) Hausser et al. [1]** | 1.6 | 1.1 | 1.8 | —— | —— |
| **(d) Lo et al. [2]** | 0.9, 0.9 | 0.8, 1.0 | 0.5, 0.1 | —— | —— |
| **(e) Scott et al. [3]** | 1.4 | —— | —— | —— | —— |
| **(f) Calabrese et al. [4]** | 1.3, 0.9 | —— | —— | —— | —— |

### B. The fitness cost per proteome fraction

To calculate the fitness cost per proteome fraction, we use the relationship between the gene copy number and the proteome fraction:

$$\phi_{\rm ex}=\frac{a_{\rm ex}}{M_{\rm tot}}p_{\rm ex}=\frac{\beta_{\rm p,ex}a_{\rm ex}}{\gamma_{\rm ex}M_{\rm tot}}m_{\rm ex}=\frac{\beta_{\rm m,ex}\tau_{\rm ex}\beta_{\rm p,ex}a_{\rm ex}}{\gamma_{\rm ex}M_{\rm tot}}g_{\rm ex}, \tag{S1}$$

where $M_{\rm tot}$ is the cellular protein mass, including the exogenous proteins. If we have two gene constructs denoted as "1" and "2", the ratio between $\phi_1/g_1$ and $\phi_2/g_2$ satisfies the following relationship:

$$\frac{\phi_2/g_2}{\phi_1/g_1} = \frac{\beta_{m,2}\tau_2\beta_{p,2}a_2/\gamma_2}{\beta_{m,1}\tau_1\beta_{p,1}a_1/\gamma_1}. \tag{S2}$$

The experimental data from Ref. [12] agree well with Eq. (S2) (Figure S1), where the two gene constructs only differ in the mRNA lifetimes.

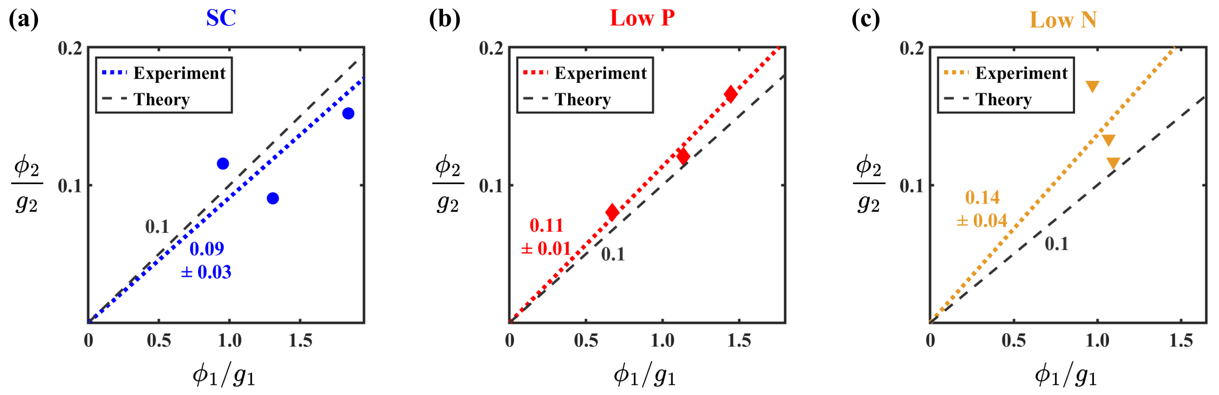


FIG. S1. **Ratio between gene copy number and proteome fraction.** The proteome fraction per gene of the wild-type and DAmP constructs (denoted as "1" and "2", respectively) in the standard culture (a), low phosphate condition (b), and low nitrogen condition (c). The mRNA lifetime of DAmP is about 0.1 that of the wild-type construct; therefore, the slope (gray dashed line) is predicted to be $\tau_2/\tau_1 = 0.1$ by (S2). The slope of the dotted line marks the experimental average of three different promoters (TDH3, PGK1, PDC1), which agrees well with our prediction.

Therefore, combining Eq. (S2) and Eq. (14) in the main text, we derive the fitness cost per excess proteome fraction as

$$\frac{C}{\phi_{\rm ex}} = \frac{\gamma_{\rm ex}M_{\rm tot}}{a_{\rm ex}}\left[\frac{1}{\tau_{\rm ex}\beta_{\rm p,ex}}\left(\frac{c_{\rm g}}{\beta_{\rm m,ex}} + \frac{T_{\rm tx,ex}}{N_{\rm n}}f_{\rm r}\right) + \frac{T_{\rm tl,ex}}{N_{\rm r}}\right], \tag{S3}$$

which is Eq. (15) in the main text. Applying $M_{\rm tot} = M/(1-\phi_{\rm ex})$, $M = g\beta_{\rm m}\tau\beta_{\rm p}a/\gamma$, $N_{\rm n}(1-f_{\rm n}) = gP_{\rm n}\Lambda_{\rm n} = g\beta_{\rm m}T_{\rm tx}$, $N_{\rm r}(1-f_{\rm r}) = mP_{\rm r}\Lambda_{\rm r} = m\beta_{\rm p}T_{\rm tl}$ (see Eqs. (1, 3, 7, 26, 27) in the main text for details) to Eq. (S3), we get

$$\frac{C}{\phi_{\rm ex}} = \frac{\gamma_{\rm ex}}{\gamma(1-\phi_{\rm ex})}\frac{g\beta_{\rm m}\tau\beta_{\rm p}a}{a_{\rm ex}}\left\{\frac{1}{\tau_{\rm ex}\beta_{\rm p,ex}}\left[\frac{c_{\rm g}}{\beta_{\rm m,ex}} + \frac{T_{\rm tl,ex}}{g\beta_{\rm m}T_{\rm tx}}(1-f_{\rm n})f_{\rm r}\right] + \frac{T_{\rm tl,ex}}{m\beta_{\rm p}T_{\rm tl}}(1-f_{\rm r})\right\}. \tag{S4}$$

Further using Eqs. (5, 13) in the main text, the fitness cost per excess proteome fraction can be expressed as

$$\frac{C}{\phi_{\rm ex}} = \frac{\gamma_{\rm ex}}{\gamma(1-\phi_{\rm ex})}\left\{\frac{\tau\beta_p}{\tau_{\rm ex}\beta_{\rm p,ex}}\left[\frac{a}{a_{\rm ex}}(1-f_{\rm t})f_{\rm n} + \frac{v_{\rm tx}}{v_{\rm tx,ex}}(1-f_{\rm n})\right]f_{\rm r} + \frac{v_{\rm tl}}{v_{\rm tl,ex}}(1-f_{\rm r})\right\}. \tag{S5}$$

The following approximations are used for Eq. (S5): (1) $a_{\rm ex}/a = L_{\rm tl,ex}/L_{\rm tl} = L_{\rm tx,ex}/L_{\rm tx}$, where we assume that the protein mass is proportional to both the length of the mRNA translated and the length of the gene transcribed; (2) $T_{\rm tx} \approx L_{\rm tx}/v_{\rm tx}$ and $T_{\rm tl} \approx L_{\rm tl}/v_{\rm tl}$ where the initiation durations are neglected because both transcription and translation primarily involve elongation as the most time-consuming process. We notice that for an "average" gene whose properties represent the average genome ($\frac{\gamma_{\rm ex}}{\gamma} = \frac{\tau_{\rm ex}\beta_{\rm p,ex}}{\tau\beta_{\rm p}} = \frac{a_{\rm ex}}{a} = \frac{v_{\rm tl,ex}}{v_{\rm tl}} = \frac{v_{\rm tx,ex}}{v_{\rm tx}} = 1$), the cost per excess proteome fraction is simplified as

$$\frac{C}{\phi_{\rm ex}} = \frac{1-f_{\rm r}+f_{\rm r}[1-f_{\rm n}+f_{\rm n}(1-f_{\rm t})]}{1-\phi_{\rm ex}} = \frac{1-f_{\rm r}f_{\rm n}f_{\rm t}}{1-\phi_{\rm ex}} \approx 1, \tag{S6}$$

where we use the approximation $\phi_{\rm ex} \ll 1$ and $f_{\rm r}f_{\rm n}f_{\rm t} \ll 1$. A numerical comparison between our model and four other models is summarized in Table S3. Intriguingly, despite distinct mechanisms behind each model, the numerical predictions for an "average" gene are close. This implies that, on average, the fitness cost due to resource competition is similar to dilution. The differences only appear when we consider a gene with distinct properties (e.g., mRNA lifetime, protein degradation rate as in Figure 4b in the main text) since a dilution mechanism (e.g., Scott et al. [3]) always gives the same $C/\phi_{\rm ex}$ for all gene constructs. The linear relationship between the fitness cost and the proteome fraction also exists in *E. coli* where the slopes are also condition-dependent (Table S4) [3, 17, 18].

TABLE S3. **A comparison of the five models on the cost per proteome fraction in both *S. cerevisiae* and *E. coli*.** Here, the numerical values represent an "average" gene in the corresponding model. We notice that an "average" gene satisfies $p_{\rm ex}/p = m_{\rm ex}/m = m'_{\rm ex}/m' = \phi_{\rm ex}$ where $m'$ is the mRNA number produced per cell cycle, such that (2) and (3) can be easily calculated from Table S1. The parameters used for (4) and (5) are (if present): *S. cerevisiae*: $\phi_{\rm Q} = 0.2$ [4], $\phi_{\rm r,0} = 0.08$, $f_{\rm r} = 0.3$ [16], $f_{\rm n} = 0.9$ [11]; *E. coli*: $\phi_{\rm Q} = 0.45$, $\phi_{\rm r,0} = 0.07$, $f_{\rm r} = 0.3$ [3], $f_{\rm n} = 0.5$ (including nonspecifically DNA-bound RNAPs which effectively enlarge the free RNAP pool [19]). The two results in *S. cerevisiae* of (5) represent the cases $n \notin$ Q and $n \in$ Q, respectively. $n \notin$ Q for *E. coli* [19]. For experimental values of various gene constructs, please refer to Figure 4b in the main text (*S. cerevisiae*) and Table S4 (*E. coli*).

| $\frac{C}{\phi_{\rm ex}}$ | **(1) This article** | **(2) Hausser et al.** | **(3) Lo et al.** | **(4) Scott et al.** | **(5) Calabrese et al.** |
|---|---|---|---|---|---|
| *S. cerevisiae* | 1.0 | 2.0 | 1.0 | 1.4 | 1.3 or 0.9 |
| *E. coli* | 1.0 | 2.0 | 1.0 | 2.1 | 2.1 |

TABLE S4. **Experimental measurements of the fitness cost per proteome fraction for *E. coli*.**

| | $C/\phi_{\rm ex}$ | Protein | Promoter | Medium | Growth Rate ($\rm h^{-1}$) | Ref |
|---|---|---|---|---|---|---|
| 1 | $1.4 \pm 0.2$ | $\beta$-lactamase | *bla* | LB | 1.29 | [17] |
| 2 | $2.0 \pm 1.0$ | $\beta$-lactamase | *bla* | M9CA | 0.99 | [17] |
| 3 | $0.9 \pm 0.6$ | $\beta$-lactamase | *bla* | M9 | 0.51 | [17] |
| 4 | $2.2 \pm 0.3$ | $\beta$-galactosidase | *lac*UV5 | M9-glycerol | 0.86 | [18] |
| 5 | $2.1 \pm 0.1$ | truncated EF-Tu | *tac* | M9-glycerol | 0.97 | [18] |
| 6 | $2.0 \pm 0.5$ | $\beta$-galactosidase | P$u$ | RDM | 1.70 | [3] |
| 7 | $1.9 \pm 0.6$ | $\beta$-galactosidase | P$u$ | cAA | 0.87 | [3] |
| 8 | $2.0 \pm 0.8$ | $\beta$-galactosidase | P$u$ | M63 | 0.60 | [3] |

### C. The extended model where the levels of endogenous proteins and the cell volume are changeable

In the main text, we mostly consider a simplified scenario in which the levels of the endogenous proteins and the cell volume are unaffected by the produced exogenous protein. In this section, we justify the simplification by taking into account all possible changes in resources ($N_{\rm r}$, $N_{\rm n}$, and $N_{\rm t}$), endogenous proteome mass ($M$), volumes ($V_{\rm c}$, $V_{\rm n}$), and mRNA lifetime ($\tau$). For a specific kind of resource $j$, where $j$ can be $r$ (ribosome), $n$ (RNAP), or $t$ (TF), by differentiating $P_{\rm j} = \frac{N_{\rm j} f_{\rm j}}{N_{\rm j} f_{\rm j} + K_{\rm j} V_{\rm j}}$ we have

$$Z_{P_{\rm j}} = (1 - P_{\rm j})(Z_{f_{\rm j}} + Z_{N_{\rm j}} - Z_{V_{\rm j}}), \tag{S7}$$

where $Z_y$ is an abbreviation for $\frac{\Delta y}{y}$ for any variable $y$ of interest. One should note that the corresponding $V_{\rm j}$ volumes for TF, RNAP, and ribosome are $V_{\rm n}$ (nuclear volume), $V_{\rm n}$, and $V_{\rm c}$ (cytoplasmic volume), respectively. All $K_{\rm j}$ are assumed to hold constant, as they are determined by the molecular binding affinities.

Differentiating the partition equation of each resource $N_{\rm j} = N_{j,{\rm f}} + N_{j,{\rm endo}} + N_{j,{\rm ex}}$ and taking $N_{j,{\rm ex}} \to 0$ (Eqs. (17, 19, 29) in the main text), where $N_{j,{\rm endo}}$ is the number of resources $j$ consumed by the endogenous genes, we get the following three equations:

$$Z_{f_{\rm t}} = \frac{1}{1-P_{\rm t}(1-f_{\rm t})}[-\frac{N_{\rm t,ex}}{N_{\rm t}} + (1-f_{\rm t})(P_{\rm t}Z_{N_{\rm t}} + (1-P_{\rm t})Z_{V_{\rm n}})], \tag{S8}$$

$$Z_{f_{\rm n}} = \frac{1}{1-P_{\rm n}(1-f_{\rm n})}[-\frac{N_{\rm n,ex}}{N_{\rm n}} + (1-f_{\rm n})(P_{\rm n}Z_{N_{\rm n}} + (1-P_{\rm n})Z_{V_{\rm n}} - Z_{P_{\rm t}})], \tag{S9}$$

$$Z_{f_{\rm r}} = \frac{1}{1-P_{\rm r}(1-f_{\rm r})}[-\frac{N_{\rm r,ex}}{N_{\rm r}} + (1-f_{\rm r})(P_{\rm r}Z_{N_{\rm r}} + (1-P_{\rm r})Z_{V_c} - (Z_{P_{\rm t}} + Z_{P_{\rm n}} + Z_{\tau}))]. \tag{S10}$$

To calculate the fitness cost $-\Delta\mu/\mu$, we differentiate Eq. (7) in the main text and get

$$Z_{\gamma} = Z_{P_{\rm t}} + Z_{P_{\rm n}} + Z_{P_{\rm r}} + Z_{\tau} - Z_{\rm M}. \tag{S11}$$

Combining Eqs. (S8-S11), approximating $\gamma \approx \mu$ since protein degradation is slow compared to cell growth, and integrating over $g_{\rm ex}$, we calculate the fitness cost to be

$$\begin{aligned} C = -\frac{\Delta\mu}{\mu} = (\frac{N_{\rm r,ex}}{N_{\rm r}} - \frac{\Delta N_{\rm r}}{N_{\rm r}} + f_{\rm r}\frac{\Delta V_{\rm c}}{V_{\rm c}})s_{\rm r} + (\frac{N_{\rm n,ex}}{N_{\rm n}} - \frac{\Delta N_{\rm n}}{N_{\rm n}} + f_{\rm n}\frac{\Delta V_{\rm n}}{V_{\rm n}})s_{\rm n}\theta_{\rm r} \\ + (\frac{N_{\rm t,ex}}{N_{\rm t}} - \frac{\Delta N_{\rm t}}{N_{\rm t}} + f_{\rm t}\frac{\Delta V_{\rm n}}{V_{\rm n}})s_{\rm t}\theta_{\rm n}\theta_{\rm r} - \theta_{\rm r}\frac{\Delta\tau}{\tau} + \frac{\Delta M}{M}, \end{aligned} \tag{S12}$$

where the symbol $\Delta$ marks a small change in the corresponding value due to the produced exogenous protein. $s_{\rm j}$ and $\theta_{\rm j}$ (j represents r, n, or t) are the sensitivity factor and the downstream factor defined in Methods A and C in the main text. The difference between this complete version and the simplified result neglecting changes in resource copy numbers and cell volume (Eq. (30) in the main text) is

$$C' = \Big(\frac{\Delta M}{M} - \frac{\Delta N_{\rm r}}{N_{\rm r}}\Big) + f_{\rm r}\Big(\frac{\Delta V_{\rm c}}{V_{\rm c}} - \frac{\Delta N_{\rm n}}{N_{\rm n}} + \frac{\Delta V_{\rm n}}{V_{\rm n}} - \frac{\Delta N_{\rm t}}{N_{\rm t}}\Big), \tag{S13}$$

where $C'$ is the cost due to protein product. The following approximations are used for Eq. (S13): $s_{\rm j} \approx 1$ and $\theta_{\rm j} \approx f_{\rm j}$ (j represents r, n, or t); $f_{\rm n} \approx 1$ [11]; the terms $-f_{\rm r}\Delta\tau/\tau$ and $f_{\rm r}f_{\rm n}f_{\rm t}\Delta V_{\rm n}/V_{\rm n}$ are neglected for simplicity, which are smaller compared to other terms and can cancel out to some extent [20], having minor impacts on the outcome.

We define $\phi'_{\rm r} = N_{\rm r}a_{\rm r}/M$ as the ribosome proteome fraction in the endogenous proteins, $\phi_{\rm n} = N_{\rm n}a_{\rm n}/M_{\rm tot}$ as the proteome fraction of RNAP, and $\phi_{\rm t} = N_{\rm t}a_{\rm t}/M_{\rm tot}$ as the proteome fraction of transcription initiation associated factors, where $a_{\rm r}$, $a_{\rm n}$, and $a_{\rm t}$ are the corresponding protein molecular masses. We introduce $\eta_{\rm nc} = V_{\rm n}/V_{\rm c}$ as the ratio of the nuclear volume to cytoplasmic volume, and $\rho = M_{\rm tot}/(V_{\rm c}+V_{\rm n})$ as the protein mass density. Then, Eq. (S13) becomes

$$C' = -\frac{\Delta\phi'_{\rm r}}{\phi'_{\rm r}} + f_{\rm r}[-\frac{\Delta\phi_{\rm n}}{\phi_{\rm n}} - \frac{\Delta\phi_{\rm t}}{\phi_{\rm t}} + \frac{\Delta\eta_{\rm nc}}{\eta_{\rm nc}} - 2\frac{\Delta\eta_{\rm nc}}{1+\eta_{\rm nc}} - 2\frac{\Delta\rho}{\rho}]. \tag{S14}$$

To derive Eq. (16) in the main text, (1) we assume a constant protein mass density such that $\frac{\Delta\rho}{\rho} = 0$; (2) we assume a constant ratio among transcriptional proteins (i.e., RNAP and TF) such that $\frac{\Delta\phi_{\rm n}}{\phi_{\rm n}} = \frac{\Delta\phi_{\rm t}}{\phi_{\rm t}} = \frac{\Delta\phi_{\rm tx}}{\phi_{\rm tx}}$ where $\phi_{\rm tx}$ is the proteome fraction of all transcriptional proteins; (3) finally, we notice $\eta_{\rm nc} \approx 0.1$ [21–24] such that $0 \approx \frac{\Delta\eta_{\rm nc}}{1+\eta_{\rm nc}} \ll \frac{\Delta\eta_{\rm nc}}{\eta_{\rm nc}}$.

---


[1] J. Hausser, A. Mayo, L. Keren, and U. Alon, Central dogma rates and the trade-off between precision and economy in gene expression, Nature communications **10**, 68 (2019).

[2] T. W. Lo, H. J. Choi, D. Huang, and P. A. Wiggins, Noise robustness and metabolic load determine the principles of central dogma regulation, Science Advances **10**, eado3095 (2024).

[3] M. Scott, C. W. Gunderson, E. M. Mateescu, Z. Zhang, and T. Hwa, Interdependence of cell growth and gene expression: origins and consequences, Science **330**, 1099 (2010).

[4] L. Calabrese, L. Ciandrini, and M. Cosentino Lagomarsino, How total mrna influences cell growth, Proceedings of the National Academy of Sciences **121**, e2400679121 (2024).

[5] M. Sette, L. A. Johnson, R. Jimenez, and F. A. Mulder, Backbone $^{1}$H, $^{15}$N and $^{13}$C resonance assignments of the 27kDa fluorescent protein mcherry, Biomolecular NMR Assignments **17**, 243 (2023).

[6] N. J. Krogan, G. Cagney, H. Yu, G. Zhong, X. Guo, A. Ignatchenko, J. Li, S. Pu, N. Datta, A. P. Tikuisis, *et al.*, Global landscape of protein complexes in the yeast saccharomyces cerevisiae, Nature **440**, 637 (2006).

[7] B. Ho, A. Baryshnikova, and G. W. Brown, Unification of protein abundance datasets yields a quantitative saccharomyces cerevisiae proteome, Cell Systems **6**, 192 (2018).

[8] G. C. Johnston, J. R. Pringle, and L. H. Hartwell, Coordination of growth with cell division in the yeast saccharomyces cerevisiae, Experimental cell research **105**, 79 (1977).

[9] B. Futcher, G. Latter, P. Monardo, C. McLaughlin, and J. Garrels, A sampling of the yeast proteome, Molecular and cellular biology **19**, 7357 (1999).

[10] F. Miura, N. Kawaguchi, M. Yoshida, C. Uematsu, K. Kito, Y. Sakaki, and T. Ito, Absolute quantification of the budding yeast transcriptome by means of competitive pcr between genomic and complementary dnas, BMC genomics **9**, 574 (2008).

[11] V. Pelechano, S. Chavez, and J. E. Perez-Ortin, A complete set of nascent transcription rates for yeast genes, PloS one **5**, e15442 (2010).

[12] M. Kafri, E. Metzl-Raz, G. Jona, and N. Barkai, The cost of protein production, Cell reports **14**, 22 (2016).

[13] C. Miller, B. Schwalb, K. Maier, D. Schulz, S. Dümcke, B. Zacher, A. Mayer, J. Sydow, L. Marcinowski, L. Dölken, *et al.*, Dynamic transcriptome analysis measures rates of mrna synthesis and decay in yeast, Molecular systems biology **7**, 458 (2011).

[14] B. Neymotin, R. Athanasiadou, and D. Gresham, Determination of in vivo rna kinetics using rate-seq, Rna **20**, 1645 (2014).

[15] P. Eser, C. Demel, K. C. Maier, B. Schwalb, N. Pirkl, D. E. Martin, P. Cramer, and A. Tresch, Periodic mrna synthesis and degradation co-operate during cell cycle gene expression, Molecular systems biology **10**, 717 (2014).

[16] E. Metzl-Raz, M. Kafri, G. Yaakov, I. Soifer, Y. Gurvich, and N. Barkai, Principles of cellular resource allocation revealed by condition-dependent proteome profiling, eLife **6**, e28034 (2017).

[17] W. E. Bentley, N. Mirjalili, D. C. Andersen, R. H. Davis, and D. S. Kompala, Plasmid-encoded protein: the principal factor in the metabolic burden associated with recombinant bacteria, Biotechnology and bioengineering **35**, 668 (1990).

[18] H. Dong, L. Nilsson, and C. G. Kurland, Gratuitous overexpression of genes in escherichia coli leads to growth inhibition and ribosome destruction, Journal of Bacteriology **177**, 1497 (1995).

[19] Y. Yan, T. Li, and J. Lin, Buffering effects of nonspecifically dna-bound rna polymerases in bacteria, Physical Review Research **6**, 033133 (2024).

[20] M. P. Swaffer, G. K. Marinov, H. Zheng, L. F. Valenzuela, C. Y. Tsui, A. W. Jones, J. Greenwood, A. Kundaje, W. J. Greenleaf, R. Reyes-Lamothe, *et al.*, Rna polymerase ii dynamics and mrna stability feedback scale mrna amounts with cell size, Cell **186**, 5254 (2023).

[21] P. Jorgensen, N. P. Edgington, B. L. Schneider, I. Rupeš, M. Tyers, and B. Futcher, The size of the nucleus increases as yeast cells grow, Molecular biology of the cell **18**, 3523 (2007).

[22] F. R. Neumann and P. Nurse, Nuclear size control in fission yeast, The Journal of cell biology **179**, 593 (2007).

[23] M. Uchida, Y. Sun, G. McDermott, C. Knoechel, M. A. Le Gros, D. Parkinson, D. G. Drubin, and C. A. Larabell, Quantitative analysis of yeast internal architecture using soft x-ray tomography, Yeast **28**, 227 (2011).

[24] Y. Wu, A. F. Pegoraro, D. A. Weitz, P. Janmey, and S. X. Sun, The correlation between cell and nucleus size is explained by an eukaryotic cell growth model, PLoS computational biology **18**, e1009400 (2022).
